# Toward naturalistic reaching: Intermediate reaches under uncertainty generalize to walking, but are not obligatory

**DOI:** 10.64898/2026.10.21.760749

**Authors:** Cady V. Seavey, Christian Seegelke, Dylan F. Cooke, Daniel S. Marigold, Tobias Heed

## Abstract

Highly-constrained laboratory-based experiments have provided important insights into movement planning. When people reach toward one of several stimuli before knowing which to target, their hand trajectories are situated intermediate to direct reaches to each stimulus. Real-world reaching, however, involves a greater decision landscape than typical laboratory tasks. We extended the go-before-you-know paradigm to a walk-to-reach task resembling unconstrained natural behavior. Participants walked three steps toward a touchscreen before reaching for one of two stimuli, marked as the target either at walk onset or after reach onset. Under uncertainty, intermediate reaches occurred, but only for some stimulus layouts and participants. Individual reaches varied considerably; many directly targeted one stimulus and corrected late. Body position at reach onset was anchored to the screen more than the stimuli and was partly intermediate. Thus, intermediate movements generalize beyond the laboratory, but as one flexible strategy among several, not as obligatory motor plan averaging.

## INTRODUCTION

Movement-related decisions are a prime example of how cognition and environment interact in complex ways in naturalistic environments: actions shape the environment, and the changing environment shapes actions (Lepora & Pezzulo, 2015; Pezzulo & Cisek, 2016). The information necessary to make movement-based decisions accumulates over time (Gold & Shadlen, 2007), and ongoing movement as well as movement choices influence which subsequent choices become available (Cisek & Kalaska, 2010; Pezzulo & Cisek, 2016). For example, when cleaning up a messy desk, new information becomes available over time as one moves and uncovers items, and this new information shapes subsequent choices of which item(s) to pick up. This continuous movement-and-decision process implies uncertainty about which movement targets may be available in the (near) future, and how to best reach them. This uncertainty, in turn, affects which movements are executed, and how.

An influential experimental paradigm for studying human reaching movements under uncertainty is the go-before-you-know task. Participants must initiate a reaching movement before they know where the movement must end. Several stimuli are shown on a monitor as potential reach targets; one of them is marked as the final target only after the participant has begun moving their hand. Reach trajectories in go-before-you-know situations lie intermediate to the trajectories participants produce when each stimulus is presented alone and there is no uncertainty about target location (Chapman et al., 2010; Gallivan & Chapman, 2014; Wispinski et al., 2018).

The occurrence of intermediate reaching movements is considered a behavioral reflection of movement uncertainty. However, there are two opposing interpretations of which mechanisms produce them. The first is that the behavior reflects an averaging of several motor plans the brain has planned in parallel (Basso & Wurtz, 1997, 1998; Chapman et al., 2010; Cisek & Kalaska, 2005; Enachescu et al., 2021; Gallivan et al., 2016; Gallivan & Chapman, 2014). Several findings support this view. First, even small differences in the stimulus layout evoke a modulation of the intermediate reach trajectory; for example, when one of several stimuli was placed slightly more left than before, the intermediate reach trajectory under target uncertainty, too, was executed slightly more to the left (Chapman et al., 2010). Second, when participants had to swiftly move, by their own choice, to one of two identical visual stimuli, early electromyographic activity reflected an influence of both stimuli (Selen et al., 2023). Third, intermediate trajectories are based on motor, not visual, information when these two aspects are disentangled using motor adaptation (Gallivan et al., 2017).

The second interpretation of intermediate trajectories is that they reflect a single motor plan that has been constructed based on the current (uncertain) context (Alhussein & Smith, 2021; Contemori & Carroll, 2025; Dekleva et al., 2018; Haith et al., 2015; Nashed et al., 2017; Wong & Haith, 2017). This notion is supported by the finding that not all uncertain situations evoke intermediate trajectories. For instance, participants used different reach strategies depending on the instructed reach speed in a go-before-you-know task (Wong & Haith, 2017): when reaches were slow, participants produced intermediate reaches; in contrast, when reaches were fast, they were directed to one of the potential targets. Likewise, when obstacles were placed such that a safe, non-collision trajectory deviated from an average trajectory, participants exhibited safe rather than intermediate trajectories (Haith et al., 2015). These strategies ensured the highest possible success rates in the respective contexts. Therefore, intermediate movements may be produced strategically to maximize task success, rather than reflecting mandatory averaging of multiple active plans.

Whereas sensorimotor decision making is often framed using natural behaviors – such as the desk cleaning example above – it is not clear whether the types of behavior identified in the lab occur in the same way in natural contexts. Studies that employed the go-before-you-know paradigm have typically implemented various constraints to tease out the described modulation of reaching movements. Some studies have participants respond quickly, with the movement completed in a narrow time interval such that trials are invalid otherwise (Alhussein & Smith, 2021; Chapman et al., 2010; Contemori & Carroll, 2025; Enachescu et al., 2021; Gallivan et al., 2016; Gallivan & Chapman, 2014; Nashed et al., 2017; Wong & Haith, 2017). Other studies have also restricted movement to a single plane of motion (Alhussein & Smith, 2021; Contemori & Carroll, 2025; Nashed et al., 2017). Such constraints thus enforce highly stereotypical movements to help uncover movement strategies. It seems plausible that the production of intermediate reach trajectories can emerge in lab experiments because the experimental context is usually narrow and overlearned, with dozens or hundreds of highly similar trials, all produced in close temporal succession.

In everyday life, reaching movements do not usually require stereotypical execution, particularly in uncertain contexts, and they often occur in tandem with full-body movement, including postural changes and walking toward the object to be handled. The decision landscape of a walk-to-reach action is considerably richer than that of typical lab-based experiments. For instance, where the foot is placed during a step affects what targets can be reached thereafter. Therefore, the hand likely starts its reach from a different location in space in each new trial, depending on where walking has positioned the body. Both the expanded decision landscape and the substantial variability of naturalistic contexts may preclude processing strategies that result in intermediate trajectories. One reason is the comparably high complexity of natural contexts. Conversely, however, participants may in fact exploit the expanded decision landscape to modulate not only reaching but also other movements: For instance, they might use walking to place their body strategically before the reach begins, similar to how participants adjust their body posture when they prepare grasping an object (Land et al., 2013; Marteniuk & Bertram, 2001). In this latter case, the laboratory-based findings about reaching movements may be viewed as a blueprint of a general, body-wide movement principle.

We addressed these questions in a go-before-you-know experiment that required participants to take several steps toward a screen on which two stimuli were displayed. As in prior, reach-only experiments, one of the stimuli changed color to mark it as the reach target. The onset of this change occurred either at the beginning of walking, so that participants were certain about the target; or it occurred after the participant had walked to the screen and begun to reach toward it, so that the reach had to be initialized under uncertainty.

## METHODS

### Participants

Twenty-two healthy adults participated in this study (4 males and 18 females; mean age = 22.6 ± 3.0 years) based on sample sizes of similar studies (Alhussein & Smith, 2021; Chapman et al., 2010; Contemori & Carroll, 2025). Participants did not have any known history of neurological, musculoskeletal, or visual diseases that could affect their walking or reaching behavior. All participants were right-hand dominant (Edinburg Handedness Inventory: 94.6 ± 9.3) and right leg dominant (based on the leg a participant would use to kick a ball). The ethics committee of the University of Salzburg approved the study protocol, and all participants gave informed written consent prior to the experiment.

### Experimental setup

Participants performed rapid reaches to a visual target. Participants performed the experiment while standing and when required to walk three steps to approach the touchscreen. We counterbalanced the order of the standing and walking blocks of trials. However, we do not include details on the standing trials because of poor data quality. Thus, we only describe the walk-to-reach protocol and report results on these trials. We displayed circular stimuli (white) and targets (green; both with diameter = 27 mm; black background) using the Psychophysics Toolbox, version 3 (Kleiner et al., 2007) on a 23” touchscreen (1920 × 1080 px; refresh rate of 60 Hz). We positioned the top of the touchscreen at 75% of the participant’s body height, measured from the floor. To track finger and walking trajectories, motion capture cameras (Visualeyez VZ4000v, Phoenix Technologies Inc., Vancouver, Canada; sampling rate 100 Hz) recorded active LED position markers placed on the right index finger (three total markers placed on the midline and medial and lateral aspects of the fingertip) and suprasternal notch. We placed additional markers on the radial styloid process, midline of the dorsal aspect of the wrist, suprasternal notch, and right and left midfeet, anterior superior iliac spines, and acromions; however, the motion tracking system did not record all positions with sufficient reliability, and we therefore used only the finger markers for our report.

### Experimental protocol

Prior to experimental trials, participants stood in front of the touchscreen and completed ten reaches toward each target without time constraints. We used their average touch location as the target location in motion capture coordinates for our data analysis. During experimental trials, the touchscreen displayed two visual stimuli, 255 mm apart (center-to-center). One of them changed its color during the trial to designate it as the reach target. This color change occurred either at walk onset (‘reach while certain’) or 50 ms after reach onset (‘reach while uncertain’), thus creating two decision contexts. Each stimulus became the final reach target with equal probability. There were two stimulus configurations; they differed only in their left-right placement on the screen, centered 88 mm to the left or right of the monitor’s center. (“stimulus set”, Figure 1B). This manipulation allows testing whether participants make adjustments of their their intermediate trajectories even to small changes of the stimulus display (Chapman et al., 2010).

**Figure 1.**
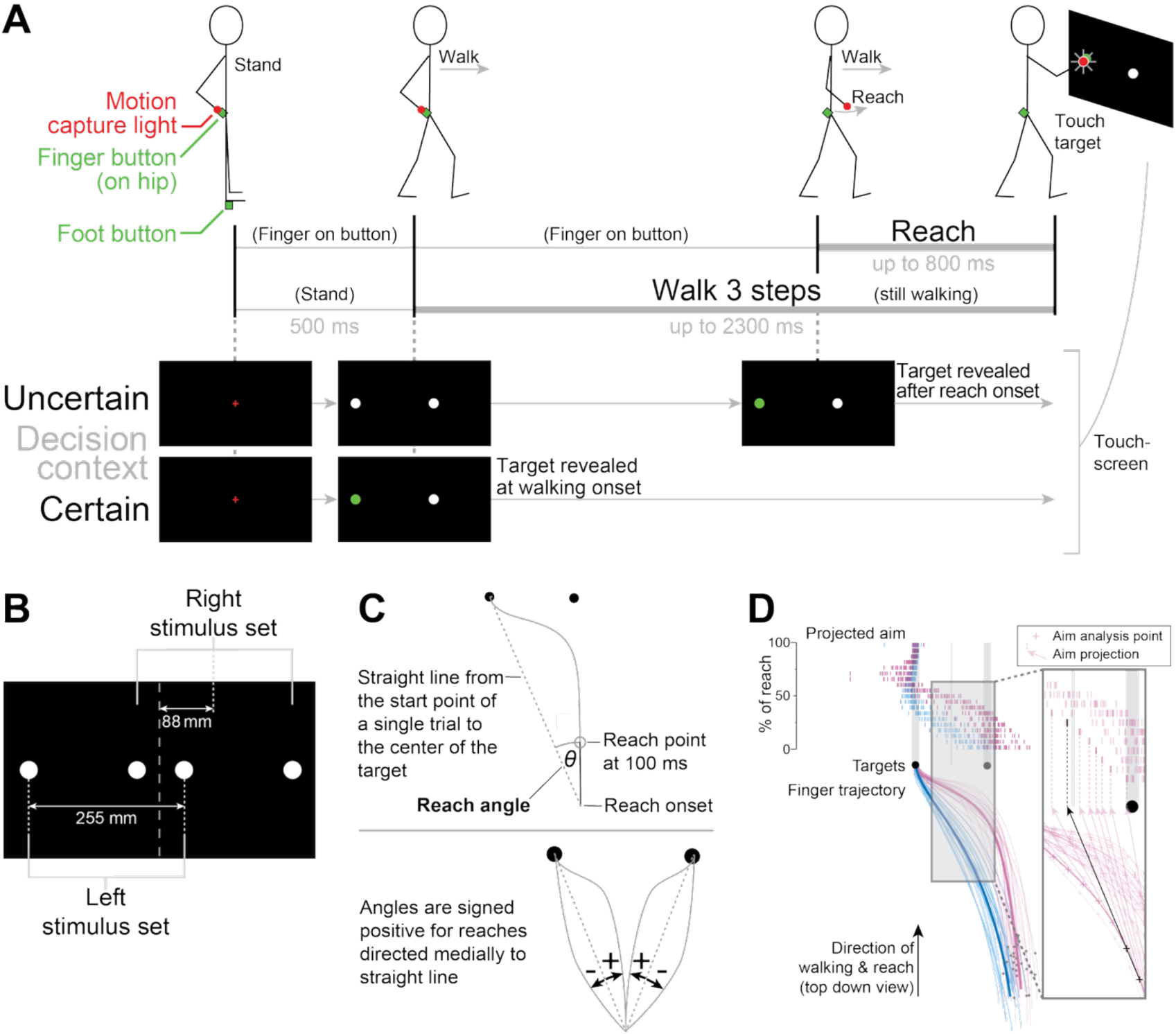
Experimental setup. (A) Participants walked toward a touchscreen initially displaying two white stimuli and reached, ultimately touching the target indicated by changing its color to green. Trials began with the participant standing, looking at the screen, and holding 2 buttons for 500 ms: their right index finger on a button on their hip and the heel of their starting foot on another button. At walk onset, the touchscreen presented the potential targets (white circles). The target was revealed (green circle) either at walking onset (“Certain” decision context) or 50 ms after reach onset (“Uncertain”). (B) There were two stimulus configurations that were shifted by 176 mm, with the stimulus midpoints shifted 88 mm to the left or right side of the touchscreen center. (C) For each trial, we calculated the reach angle between a straight line from the reach onset to the center of the target and the reach point 100 ms along the trajectory (top). Reach angle was signed positive for reaches directed medial to a straight line to the target (bottom). (D) Aim analysis (inset) with example reach trajectories and progression of reach aim. Every 5% along each reach trajectory (+, aim analysis points), we projected the aim to the touchscreen. Rasters above indicate left/right position of the aim. Each raster row indicates the progression of the aims across trials in increments of 5% of the reach. Inset shows expanded view of boxed area with one example reach trial trajectory highlighted and annotated to show how aim rasters were determined. Aim lines with arrowheads extend from consecutive aim analysis points from the example trajectory and end at the anterior-posterior position of the targets. This left/right position is projected vertically (dashed lines). Later aim rasters are plotted progressively higher up. A single example aim line and raster are plotted in black. Additional description in Figure 4.

Participants began each trial standing at a marked location about three steps away from the touchscreen and were required to depress two buttons for 500 ms: Their right index finger was placed on a hip button, and the heel of their preferred starting foot was placed on a button located on the ground. At walk onset, the touchscreen presented the potential targets. To encourage rapid reaching, we constrained the total walking time to 2300 ms (foot button release to finger-screen contact) and the total reach time (which occurred during walking) to 800 ms (hip button release to finger-screen contact). At the end of each trial, the touchscreen indicated whether the participant had met these timing constraints. Our task consisted of two target decision contexts (‘reach while certain/uncertain’), two stimulus midpoints (stimulus sets shifted left and right), and two endpoints (target endpoint: left and right target of a stimulus set) for a total of eight conditions. Participants completed 25 trials per condition that met timing constraints for a total of 200 trials. To avoid fatigue, we divided the experiment into five blocks. Each block consisted of five repetitions of each condition, randomized within the block for a total of 40 reaches. If the participant did not meet timing constraints for a given trial, that trial was repeated at the end the block. Participants could rest between blocks. Prior to starting the task, participants completed one practice block of trials.

### Data and statistical analysis

We analyzed kinematic data using custom code written in MATLAB. We interpolated missing data for gaps under 16 frames using a cubic spline; we left larger gaps marked as missing data. Then we filtered the data using a zero-lag, fourth-order, low-pass Butterworth filter with a cutoff frequency of 6 Hz. We defined walk onset as the time of release of the foot button, and reach onset as the time of release of the hip button. We defined reach endpoint as the time the touchscreen registered the participant’s touch. We removed five participants from the analysis due to poor data quality (< 5 useable trials for all conditions; these data are, however, included in the available dataset). From remaining participants, we excluded trials if the frame at reach onset and/or the data point at 100 ms after reach onset was missing after interpolation (31% of trials were excluded for this reason), or if the participant touched the wrong side of the screen (< 1%). For one of the four target locations, three participants contained < 5 useable trials for one decision context (reach while certain/uncertain) but not the other. In these cases, we removed the participant’s data of both decision contexts for the respective target from analysis. Subsequent analyses thus included data from 17 participants, three of which had some target locations excluded. For all statistical analyses, we used JMP 18 software (SAS Institute Inc., Cary, NC). We considered effects significant at an alpha level of 0.05. We used Tukey’s post hoc tests to follow up significant interactions.

We asked whether participants’ initial reach strategy depended on the decision context. We expected participants to reach between the two stimuli when initiating their reach before knowing which stimulus was the target (“uncertain”). In contrast, we expected a direct reach to the target when it was already known at the beginning of the trial (“certain”). We used the initial reach angle as a measure of reach direction at the beginning of the reach. Initial reach direction indicates the feedforward motor plan prior to any online corrections, because sensory information is not yet available due to neural transmission times (Scott, 2016). We defined initial reach angle as the angle between a straight line from the fingertip position at reach onset to the target and the location of the fingertip marker 100 ms after reach onset, calculated using the inverse tangent function. We designated angles directed medial of the straight line as positive (i.e., clockwise angles for left targets, counterclockwise angles for right targets; see Figure 1C). We tested differences in initial reach angle between factors decision context (reach while certain vs. uncertain), target endpoint (left vs. right), and stimulus set (sets shifted left vs. right) with a three-way linear mixed-effects model, and using participant as a random effect.

While initial reach angle quantified the deviation of the initial aim from a straight-line trajectory to the target, it was not informative about whether a participant aimed at the same location across trials because the starting position varied between reaches. Thus, we performed an aim analysis. We first time normalized the data and then identified the position of the fingertip along the reach trajectory at every 5 percent of the reach time to determine the location of the aim. We constructed a line between consecutive points along the trajectory and extrapolated the line to the anterior-posterior position of the targets. We indicated the projected aim for each trajectory as vertically dash lines and represented time upward across rows (Figure 1D). This analysis assesses how aim evolved over the duration of the reach and whether participants adopted different reaching strategies in the different decision-context conditions. Subsequently, we averaged trials for each condition within participants and also computed group averages.

Participants did not receive specific instructions about how or where they had to walk beyond the general instruction to take three steps and reach for the target. Therefore, participants could theoretically position their body strategically relative to the screen or stimuli prior to reach onset. Participants held their pointing finger on the hip button until they began their reach. We used finger position prior to hip button release as a proxy of body position in space and identified the left/right fingertip position at reach onset relative to the center of the touchscreen. We analyzed the left/right position of the finger at reach onset with a three-way linear mixed-effect model with fixed factors decision context (reach while certain vs. uncertain), target endpoint (left vs. right), and stimulus set (sets shifted left vs. right), and participant as a random effect.

## RESULTS

### Trajectories reflect the unresolved target options

We first asked whether reach trajectories are adjusted based on (un)certainty, even when a person walks toward the potential targets. Reach trajectories were mostly straight and directed toward the target when the target had been specified at walk onset (reaching while certain, Figure 2). In contrast, when the target was specified only after the reach had been initiated (reaching while uncertain, Figure 2), reach trajectories often curved toward the target after it was cued. Whereas this finding is, at first glance, compatible with the previous reports of intermediate reach trajectories (Chapman et al., 2010; Gallivan & Chapman, 2014), trajectories were not all intermediate under uncertainty: In some cases, participants produced trajectories akin to those observed in previous go-before-you-know experiments that lie between the direct reaches initiated with certainty about target location (e.g., Figure 2, Participants 8 and 14, right column). In other cases the trajectory was initially directed toward one of the two stimuli. If the opposite stimulus became the target, the reach turned at the very end toward the correct stimulus (e.g., Figure 2, Participant 17). Not all cases are easily classifiable. For example, for both Participants 8 and 14, uncertain reaches toward the left stimulus set were strongly biased toward the right target, but the average trajectory nonetheless shows a slight leftward bias visually. The same is true for the average of the full sample (Figure 5, left side).

**Figure 2.**
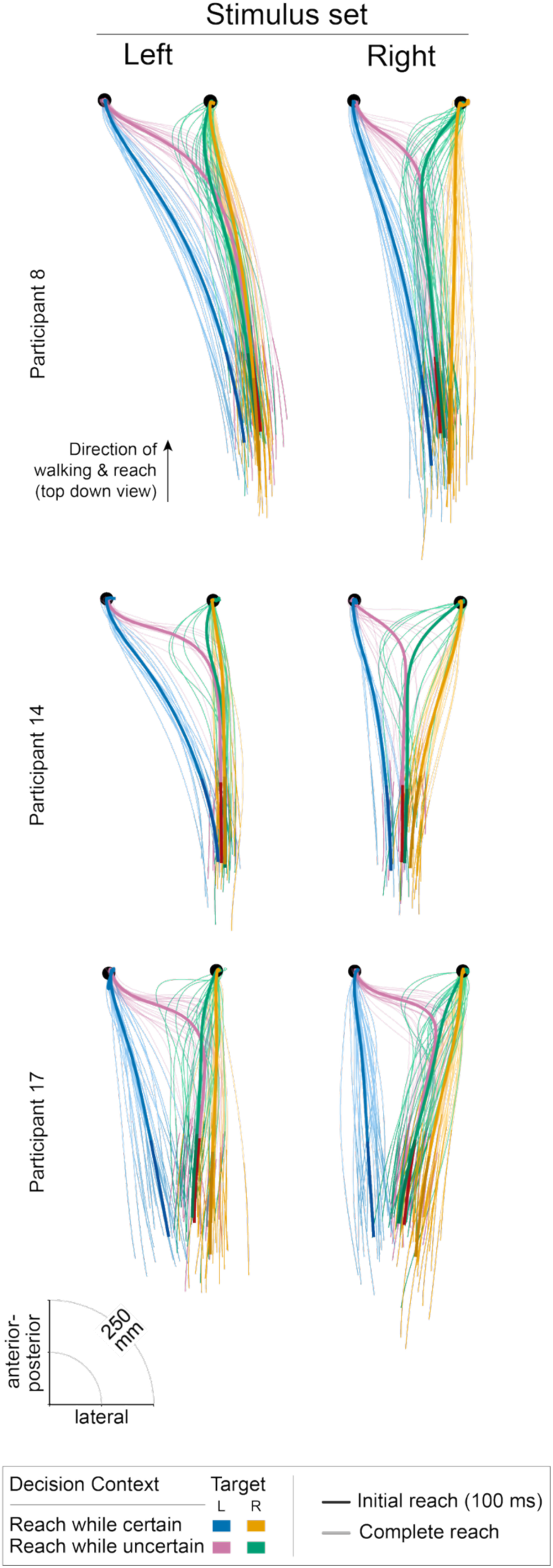
Target (un)certainty and location influence reach trajectory. Example reach trajectories from three participants. Blue and orange traces: reach while certain. pink and green traces: reach while uncertain. Bold lines indicate condition-wise average trajectories, and fine lines indicate individual trajectories. Darker hues indicate the first 100 ms of the trajectory.

Initial reach direction differed depending on decision context. When reaching while certain, participants directed their initial reach toward the cued target (Figure 2, blue and orange colors). When reaching while uncertain, some participants directed their initial reach more toward the center of both potential targets (e.g., Participants 8 and 14, Figure 2, pink and green colors) when presented with the right stimulus set. However, when presented with the left stimulus set, participants generally biased the direction of their reach toward the right stimulus. Other participants (e.g., Participant 17, Figure 2) biased the direction of their reach toward the right stimulus irrespective of the stimulus set. To quantify this behavior, we calculated initial reach angle (see Figure 1C). Initial reach angle differed based on decision context, target endpoint, and stimulus set (three-way interaction: F_1,103_ = 7.43, p = 0.008). To interpret this interaction, we evaluated the decision context × target endpoint interaction for the left and right stimulus sets separately. For the left stimulus set, reaching while uncertain increased the initial reach angle compared to reaching while certain for the left but not for the right target (Figure 3A, left side; Figure 3B, left side; decision context x target endpoint interaction: F_1,39_ = 37.08, p < 0.001). This is also evident in the examples shown in Figure 2. For the right stimulus set, reaching while uncertain increased initial reach angle compared to reaching while certain for both the left and right targets (Figure 3A, right side; Figure 3B, right side; F_1,47_ = 11.31, p = 0.002). This result indicates decision context affected initial reach direction for both targets (see also examples in Figure 2, Participants 8 and 14).

**Figure 3.**
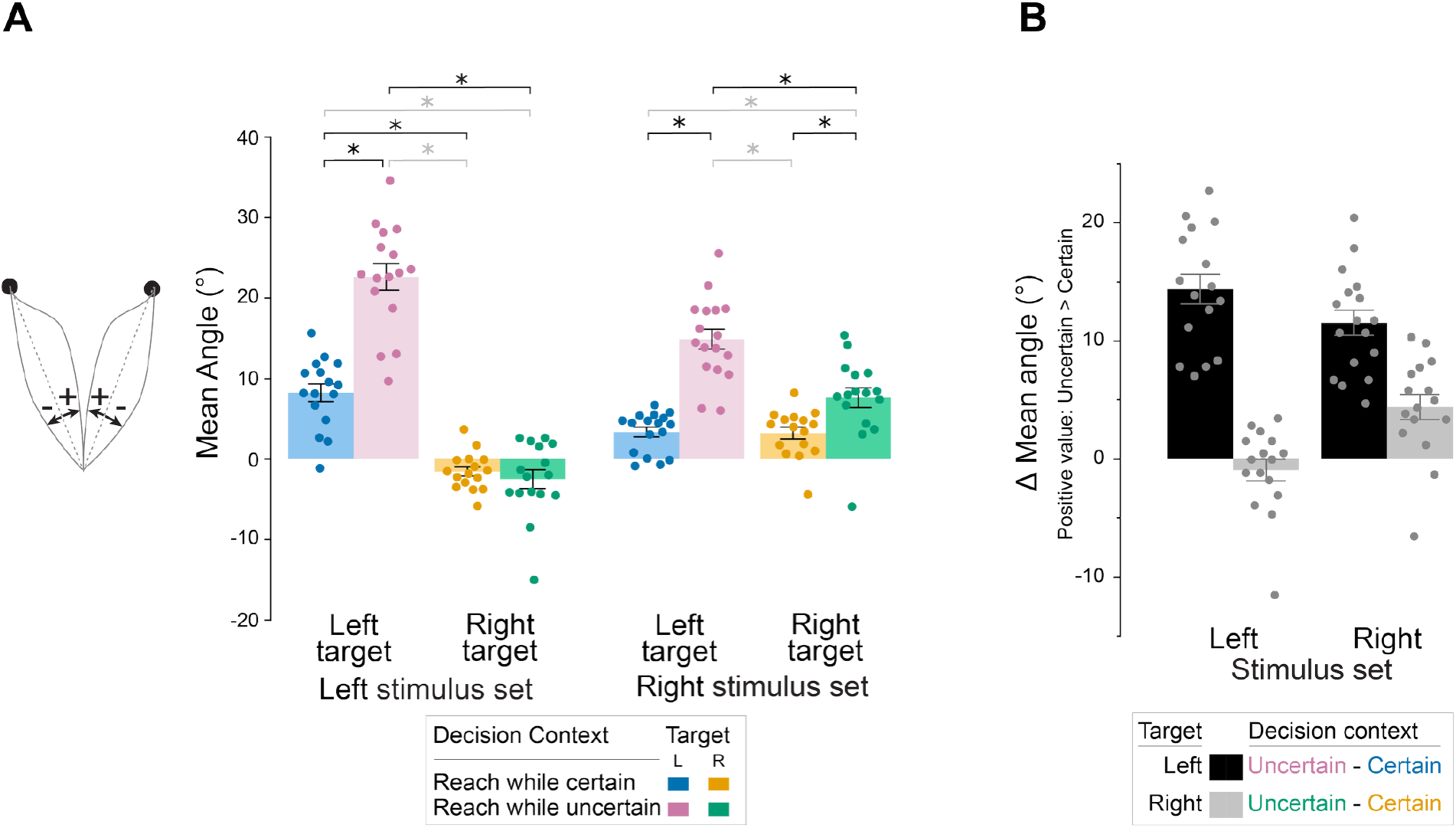
Effect of uncertainty on initial reach direction. (A) Mean ± SE initial reach angle across participants (n=17). *Significantly different based on post hoc tests after significant decision context x target endpoint interaction (*P* < 0.05). Black * and brackets, significant comparisons of primary interest. (B) Mean ± SE difference in initial reach angle between reaching while certain vs. uncertain. Positive values indicate greater initial reach angle when reaching while uncertain. The means for each participant are superimposed.

### Initial aim strategy is highly variable: precise aiming does not occur until late into the reach

Previous research used the initial reach angle to characterize initial reach bias (Contemori & Carroll, 2025), just as we did in the present study. Here, with each reach starting at a different location, the direction to the target, too, differs in each trial. Our initial reach angle measure compensates for this potential confound because it is computed relative to a straight line to the target (see Figure 1C). However, by removing the relevance of the starting location, the measure removes the absolute information about where the reach was directed. In particular, if a participant started their reach more toward the left versus right on a trial, analyzing (non-zero) initial angles alone would not tell us whether the participant was aiming at a common location across trials. Moreover, focusing on the initial reach angle disregards whether and how the aim evolves throughout the reach. Therefore, we computed the reach aim at different points along the reach (see Figure 4) and extended it to the frontal plane on which the potential targets lie.

**Figure 4.**
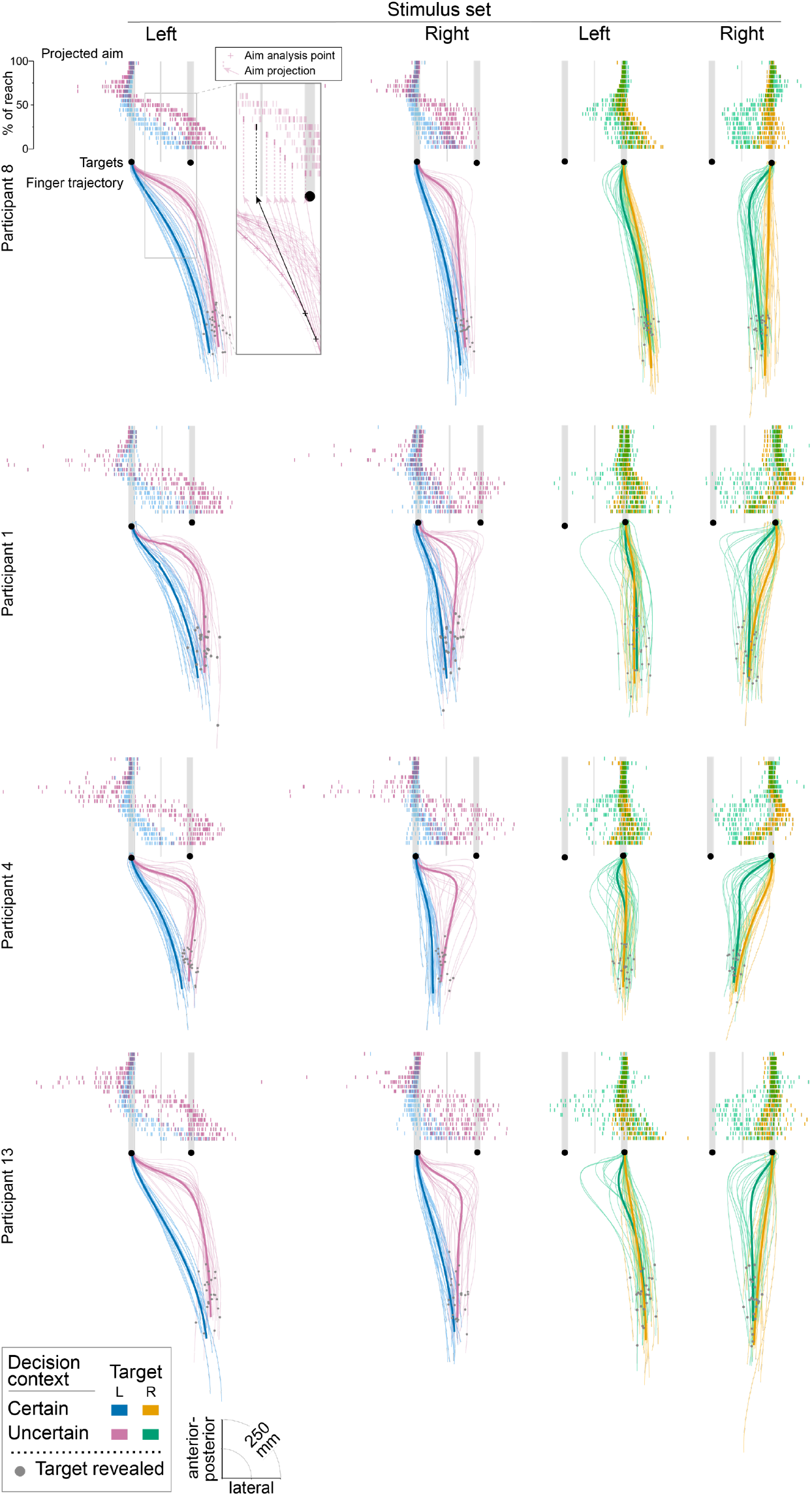
Reach aim progression varies within and across participants. Progression of reach aim over time for four example participants. Fine trajectories are individual, time-normalized trials. Rasters above indicate left/right position of the aim projected to the anterior-posterior position plane of the targets. Each raster row indicates the progression of the aims across trials in increments of 5% of the reach with later aim rasters plotted progressively higher up. See Figure 1 for details of aim analysis shown in in-set.

Initial aim strategy was highly variable within and between participants. When reaching with uncertainty, some participants aimed toward the midpoint of the two potential targets (Figure 4, Participant 8, right stimulus set, both targets: pink and green rasters). However, when reaching with uncertainty toward the potential targets with the other stimulus set, the same participant aimed toward one of the potential targets (Figure 4, Participant 8, left stimulus set, both targets: pink and green rasters). These differences between the two stimulus sets thus mirror the results of the initial reach angle analysis. Yet, even within the same condition, participants varied their reach behavior from trial to trial, suggesting individuals did not aim consistently. This is demonstrated by the spread of the aim rasters spanning the width of the potential targets (e.g., Figure 4; all Participants, right stimulus set, left target: pink rasters). This can also be seen in the wide spread of pink trajectories in all Figure 4 participants. Indeed, Participant 1 appears to have a bimodal distribution of trajectories in this condition. Aiming also varied between participants. The spread of rasters for the initial aim in the group figure demonstrates variance in where participants each aimed on average (Figure 5).

**Figure 5.**
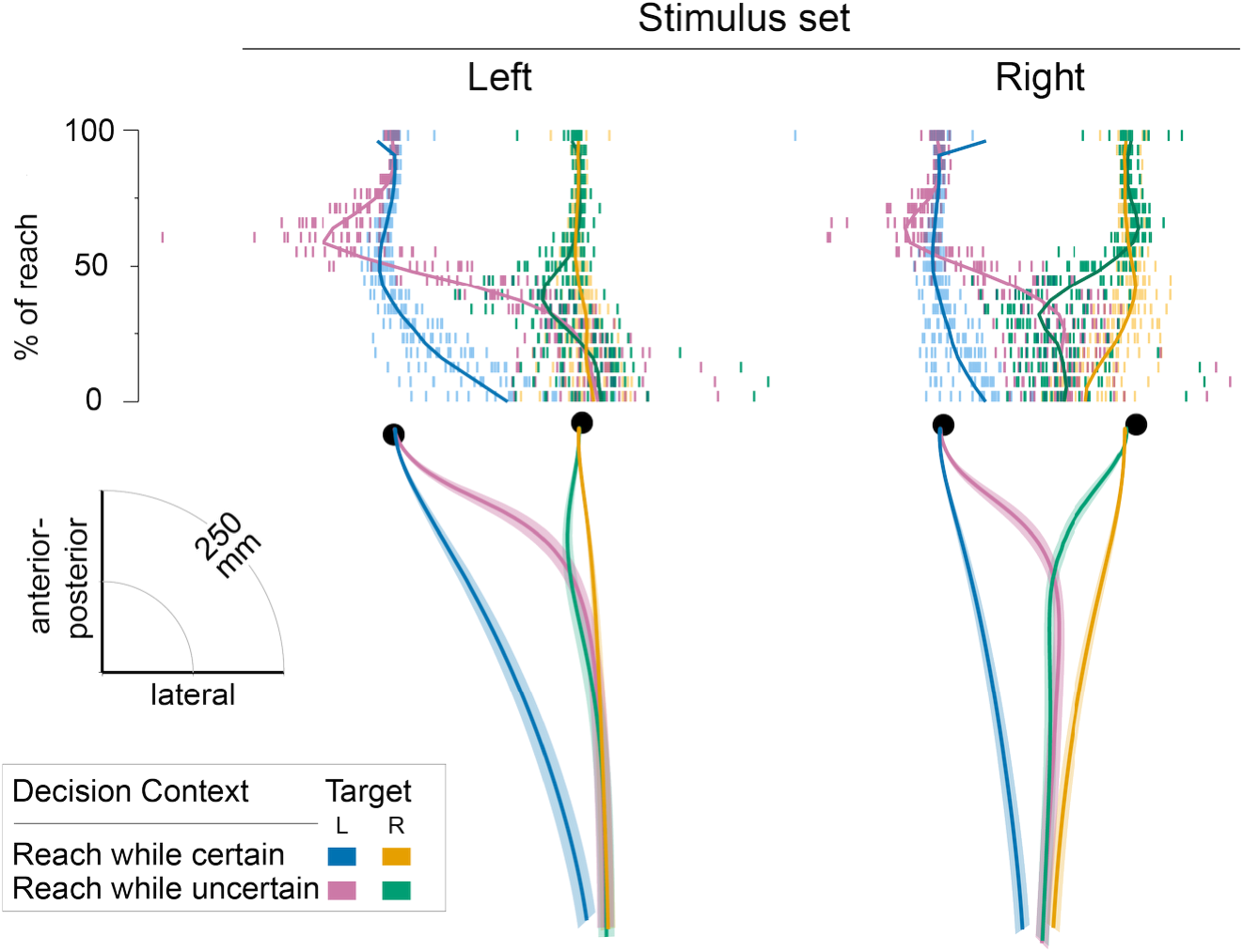
Average reach trajectories and aims. Each raster row indicates the average progression of the aims in increments of 5% of the reach. Each raster of a given color in a row is the mean aim of one participant at that timepoint. The trace superimposed on the raster plots indicate condition-wise group average aim. Lines and shading below indicate condition-wise group average trajectories and 2D SE error bands, respectively. Error bands can be conceived of as exactly enclosing the series of standard error ellipses (xSE, ySE) centered on each datapoint along the path.

### Target (un)certainty modulates positioning before reach onset

Previous lab-based studies enforced a very precise, fixed starting position for the finger. In our experiment, however, walking toward the screen made the finger’s starting position variable. In fact, participants may have positioned their body—and accordingly, the hand’s starting position— strategically to reduce the motor cost of reaching toward the target. For instance, when reaching with uncertainty, they might have positioned their body (and thus, fingertip) between the visual stimuli. Consequently, participants may have reduced the overall minimal cost of reaching to either stimulus. In other words, it would have been possible for participants not only to adjust their reach (as shown above and in previous experiments), but also its starting point (through strategic body placement).

The example trajectories illustrated in Figure 2 show that, regardless of decision context, the fingertip’s (and hip’s) starting location varied across conditions. Specifically, finger starting position was often between the two visual stimuli when they were presented toward the right side of the touchscreen (right stimulus set; see e.g., Figure 2, all depicted participants, right column).

In contrast, some participants positioned their fingertip more in front of the right visual stimulus when the stimuli were presented toward the left side of the touchscreen (left stimulus set, see e.g., Figure 2, Participants 8 and 14, left column). Thus, those participants did not position their body based on just the stimuli, but probably used other scene characteristics, such as the position of the touchscreen, to place themselves.

At the group level, participants placed their body in front of the visual stimuli prior to reach onset differently depending on the experimental condition. We determined left/right finger starting position relative to the left/right center of the touchscreen. Thus, if participants placed their body only relative to the monitor as a landmark, we would expect no difference in body position for the two stimulus sets, and if participants tried to position themselves in the middle of the monitor, start positions should be near zero. However, finger starting position differed significantly between stimulus sets (Figure 6; main effect: F_1,106_ = 105.22, p < 0.001). Specifically, finger starting position relative to touchscreen center for the left and right stimulus sets was 73.7 ± 48.0 mm and 97.7 ± 40.0 mm, respectively; thus, the finger started in front of the right half of the monitor for both stimulus sets, but 24 mm further to the left for the left stimulus set.

**Figure 6.**
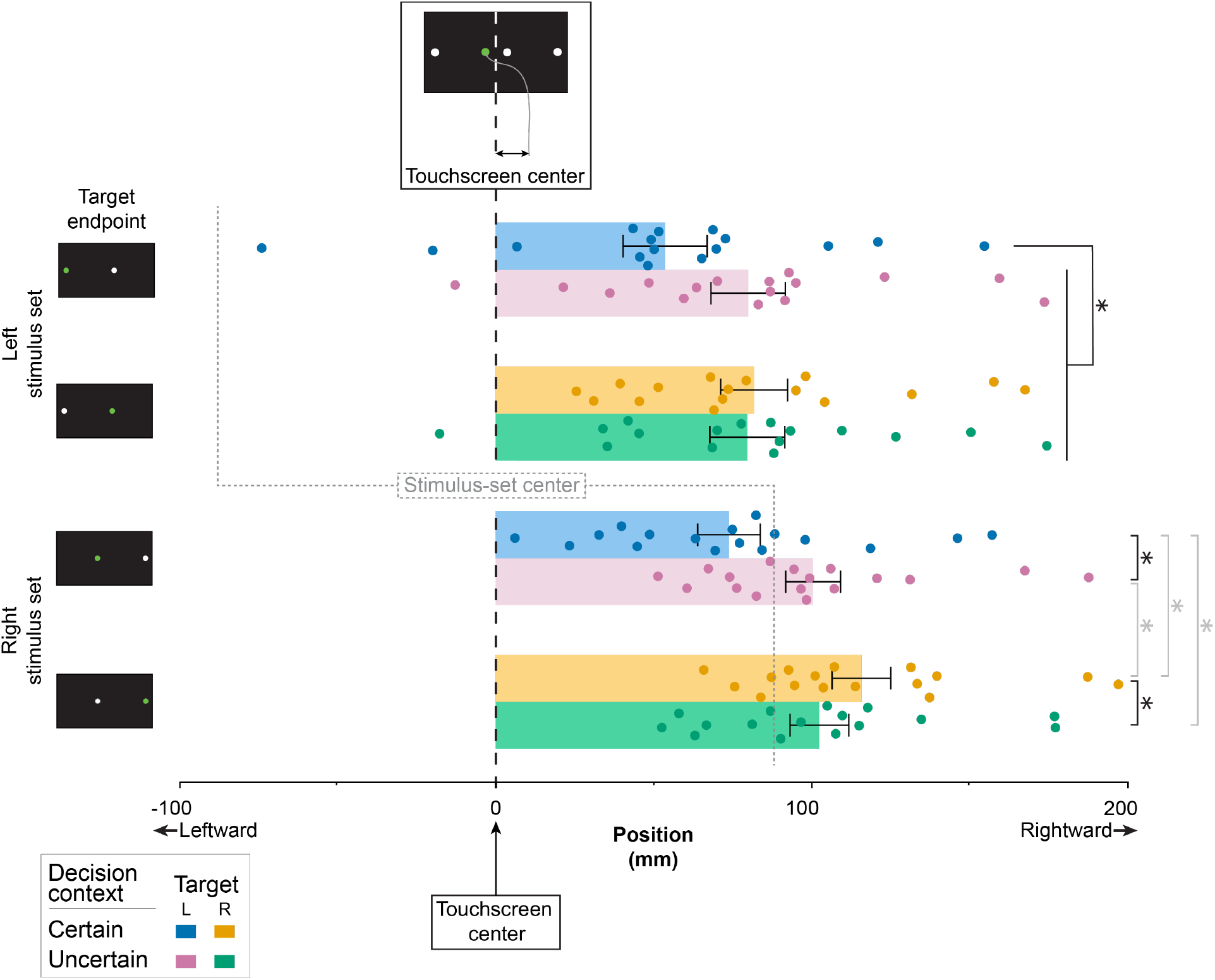
Effect of stimulus set, decision context, and target endpoint on left/right finger position at reach onset Left/right finger starting position relative to touchscreen center at reach onset across participants (mean ± SE, n=17). The means of each participant are superimposed. Dotted gray lines indicate stimulus center for each stimulus set, illustrating that participants started their reaches overall biased to the right also for the left stimulus set. The black dashed line indicates the touchscreen center, illustrating that reaches began on the right side of the screen for both stimulus sets. * After separating left and right stimulus sets, post hoc tests for significant decision context x target endpoint interactions revealed differences in finger starting position between several conditions (*P* < 0.05). Black * and brackets, significant comparisons of primary interest.

In addition, finger starting position depended on decision context and target endpoint (interaction of decision context x target endpoint: F_1,106_ = 51.0, p < 0.001). As is evident in Figure 6, the finger’s starting position differed for the left and right target when participants knew they had to reach these locations (orange vs. blue difference of 28.2 mm and 41.9 mm for the left and right stimulus sets, respectively). In contrast, finger starting position was indistinguishable for left and right targets under uncertainty—as would be expected, because the target wasn’t known at reach start (green vs. pink difference of 0.2 mm and 1.9 mm for the left and right stimulus sets, respectively).

However, we had additionally expected that participants would place their finger intermediately in the uncertain relative to the certain decision context, in analogy to the modulation of reach trajectories. Both Fig. 2 and Fig. 6 suggested that participants indeed showed this behavior, but only for the right stimulus set—just like intermediate trajectories had been evident primarily for the right stimulus set. Therefore, we analyzed the data of the left and right stimulus sets separately, although interactions with stimulus set were not significant (3-way interaction: F_1,106_ = 1.45, p = 0.232; stimulus set x decision context: F_1,106_ = 1.27, p = 0.262; stimulus set x target endpoint: F_1,106_ = 0.51, p = 0.475). This analysis confirmed the visual result pattern.

For the left stimulus set, finger starting position differed depending on decision context and target endpoint (F_1,44_ = 25.28, p < 0.001). Post hoc tests revealed that finger starting position for certain left-target reaches (Figure 6, top half; blue bar) differed significantly from all others (p < 0.001), whereas those did not differ significantly from each other (p = 0.826 – 0.988). Thus, participants positioned their finger under uncertainty in the location they also placed it when reaching directly toward the right target. This result pattern resembles that of the reach trajectories (see Figures 2 and 5) and does not indicate intermediate finger placement when uncertain.

For the right stimulus set, the decision context x target endpoint interaction was again significant (F_1,46_ = 74.95, p < 0.001). Post hoc tests revealed a significant difference in finger starting position between decision contexts for both the left (p < 0.001; blue vs. pink, lower half of Figure 6) and right (p = 0.001; orange vs. green) targets, whereas the finger starting position did not differ between the uncertain reaches that later had to be directed to the left and right target (p = 0.997; pink vs. green). Thus, the finger starting position was intermediate for uncertain reaches.

Taken together, both touchscreen and stimulus-set locations affect finger starting position. Overall, these results suggest that movement adjustment is not confined to hand trajectories but rather extends to other body parts and movement types as well as to the movement that precedes the actual reach.

## DISCUSSION

We expanded the well-known go-before-you-know reaching paradigm by requiring participants to approach the stimulus-displaying touchscreen with several walking steps. This manipulation increases the decision landscape of the task and the variability of multiple movement parameters, such as body posture and the hand’s starting position, and frames the experiment in a natural walk-to-reach context. Previous research showed that in an uncertain, go-before-you-know decision context, people initiate reaches intermediate to direct trajectories to potential targets. (Chapman et al., 2010; Gallivan & Chapman, 2014; Wispinski et al., 2018).

Our experiment revealed three key findings. First, intermediate reach trajectories occurred when participants walked toward the screen and the target was only revealed after reach onset. Thus, intermediate reaching under uncertainty occurs in a context that is less constrained than typical sensorimotor laboratory experiments. Second, the decision context affected body placement at reach onset, measured as the finger’s starting position on the hip. This effect was small compared to both variability and the size of the stimulus set shift. Third, in contrast to previous work using this paradigm, we observed variable walking/reaching behavior across specific stimulus configurations and across and within participants. Some variability may be related to the increased decision landscape of our experimental design compared to reaching that is isolated and highly stereotypical, indicating that the expanded task context modulated how trajectories are planned. Variability may have also been present in prior work but hidden by averaging. Inter-and intra-participant variability may imply that participants chose among several strategies. Below, we further discuss and integrate these three findings.

Laboratory experiments in sensorimotor neuroscience often place strong constraints on how participants may execute their movements. This approach allows for teasing out small and difficult-to-observe systematic movement modulations. Yet, laboratory designs may not generalize to real-world contexts: they may fail to elicit cognitive processes and decision-making strategies that are relevant in real-world settings, or, conversely, induce processing strategies that are rarely employed outside the laboratory (Cisek & Green, 2024; Gordon et al., 2021; Ingram & Wolpert, 2011; Krakauer et al., 2017; Maselli et al., 2023; Nastase et al., 2020). Our experiment demonstrates that the go-before-you-know intermediate trajectory strategies identified in isolated reaching experiments do generalize to far less constrained, natural walk-to-reach behavior. Our experiment, therefore, principally validates the strict methodology applied in previous work for more complex and closer-to-real-life contexts.

At the same time, the modulation of movement with action (un)certainty was not restricted to arm reaches. Because participants rested the reach hand’s finger on a button attached to the hip before each reach, we could use finger position during the walking phase as a proxy for hip or body position. The finger’s (and with it, the body’s) placement differed for the left and right stimulus sets as well as for the decision context. However, these adjustments were small. Body placement remained closely aligned to the monitor (or the room overall): participants only compensated for roughly 14% of the spatial displacement between the two stimulus sets (24 mm of the 176-mm shift). Distributed movement modulation across the entire body would be in line with theoretical proposals of continuous sensorimotor decision-making (Gordon et al., 2021; Lepora & Pezzulo, 2015; Pezzulo & Cisek, 2016) as well as balance recovery responses involving coordinated muscle activity across body parts following perturbations (Lowrey et al., 2017; Marigold et al., 2003; Pijnappels et al., 2010). Moreover, previous walk-to-grasp experiments suggest that participants trade off between walking and reaching depending on their relative cost or effort (Potts et al., 2018; Rosenbaum, 2008). Therefore, body placement might exhibit clearer preparatory effects if some aspects of our walk-to-reach task were adjusted—for instance, if no screen edge were visible that could serve as reference frame for the walking approach.

Our findings are in conflict with the idea that reach trajectories reflect average movements derived from direct reaches to the various target options (cf. Chapman et al., 2010). Such averaging cannot explain why intermediate trajectories in our study were pronounced for only one stimulus set. Moreover, if intermediate reach trajectories reflected obligatory averages of individual reach plans, then all trajectories of a given condition should be comparatively similar. However, within a given participant and condition, single trajectories were highly variable and not aimed consistently at a hypothetical target between the two stimuli of a stimulus set. Rather, the present results may fit two alternative interpretations.

The first interpretation is that uncertain trajectories are all produced under one common reach strategy that involves no early averaging of movements. This strategy entails coarsely propelling the hand forward in the general direction of the stimuli, and then performing a late, visually-guided correction toward the instructed target. In this propel-and-correct view, the largely stereotypical placement of the body and hand at reach start and the high variability of individual reaches suggest that participants do not even try to prepare, because the reach target is revealed so late. Our experiment implemented a time constraint to avoid participants simply stopping their movement to wait for the target to be revealed. Although the time constraint was twice as long as the original reach-only study (Chapman et al., 2010), participants may still have judged it too short in our more complex task context to use detailed planning. As a consequence, biomechanical constraints— which differed for the left-and right-shifted stimulus sets because participants always began their reaches on the right side of the screen—and the variability of the hand’s start point may have largely determined the trajectory before the final corrective movement. Moreover, the average-like trajectories observed for the right stimulus set may have occurred because participants could simply propel the hand forward from their preferred, right-side starting point, to guide the hand in the general direction of the stimuli. Given that the hand began the reach from the right side of the body, we would expect propelled reaches to be biased toward right side of the left stimulus set. This interpretation may also explain why participants did not exploit the possibility of positioning their body strategically: the uncertainty inherent in the task may have induced late-correction strategies, which in turn may have led to discounting fine-grained preparatory positioning.

In the second interpretation, people make average movements to potential targets but do so only sometimes, and/or other factors obscure or cancel out the intermediate strategy. People may choose different reaching strategies based on individual preferences and contextual factors. Various findings point toward the availability of multiple strategies rather than an obligatory integration of all potential targets into an intermediate trajectory. For instance, one study reported that participants sometimes appeared to initiate reaches directed to one stimulus, though they more often produced intermediate trajectories (Contemori & Carroll, 2025). Others have observed more switching than intermediate reaching when the motor cost for switching was low (Burk et al., 2014), again suggesting weighing of multiple decision aspects. Finally, participants produced stimulus-directed rather than intermediate movements when stimuli were so far apart that they could not be reached by late correction in the given time constraint (Wong & Haith, 2017), and when a non-intermediate trajectory avoided collision with an obstacle (Alhussein & Smith, 2021). Thus, in our task, uncertain reaches directed toward one stimulus may have reflected, after choosing for that trial, a late-correction strategy; intermediate reaches may, in contrast, have reflected integration of all target locations for that trial (Chapman et al., 2010; Gallivan et al., 2017; Selen et al., 2023). Alternatively, movement averaging may always take place, but other factors like biomechanical constraints (e.g., contralateral—or cross-body—reaches, more on this below) may erase behavioral evidence of this computation.

Overall, we lean more strongly toward the second interpretation, which allows for strategic use of averaging depending on preference and context. Participants only propelling their hand (our suggested first interpretation) cannot explain the intermediate positioning of the hand’s starting position. Moreover, many individual trajectories of the uncertain conditions project clearly between those of the direct reaches to the left and right targets. These factors make it unlikely that participants exclusively used a propel-and-correct strategy.

Our experiment did not disentangle which concrete aspects participants might integrate into their decisions. However, it seems likely that at least the precise position of the last step before the reach and biomechanical constraints that emerge between body position and target locations play a role. For instance, ipsilateral reaching requires an elbow extension, whereas cross-body reaches typically require elbow flexion, trunk rotation, and shoulder movement (cf. Gallivan & Chapman, 2014). Thus, participants may have prioritized the ipsilateral reach to the right target in the uncertain decision context because of the resulting lower motor cost. Moreover, participants may have perceived contralateral reaches as more effortful: longer movement duration is associated with higher perceived effort (Morel et al., 2017), and contralateral reaches typically have a longer duration (Carey et al., 1996). Others have argued that it is the transformation from visual to proprioceptive space that induces reach biases (Wang et al., 2026). Finally, the task context, such as pacing and time constraints, likely affects strategic choices. Thus, multiple factors probably shape each single reach in concert.

In conclusion, people make intermediate reaches when faced with multiple potential target locations even when the decision landscape is expanded. This extends findings from highly constrained laboratory settings that isolate and standardize reaches to more naturalistic behaviors. However, our naturalistic extension of the paradigm revealed higher variability between participants and stimulus sets than implied by previous, highly constrained reaching-only tasks. This finding suggests that the higher complexity of the walk-to-reach task’s decision landscape prompted participants to use various decision strategies. Moreover, reaching was shaped by additional aspects such as scene characteristics and the spatial characteristics of the stimulus set. Intermediate reaching, therefore, appears to be strategically chosen rather than obligatory. Another strategy may have been to rely on late, visually guided correction of generic reach movements. However, intermediate pre-reach positioning of the body and hand argues against the propel-and-correct interpretation as a dominant or exclusive strategy participants applied throughout. In sum, these findings lead us to favor a nuanced interpretation: that intermediate reaches are sometimes deployed strategically among several possible approaches, explaining their absence in some contexts and trials. From a methodological viewpoint, the rich behavioral variability makes the walk-to-reach approach an excellent candidate paradigm for further exploration of movement decisions in dynamic, real-world environments.

## ACKNOWLEDGMENTS

We thank Juliana Möller and Paul Camp for their help with data acquisition.

This study was supported by an Alliance International Catalyst grant from the Natural Sciences and Engineering Research Council (NSERC) of Canada (Grant number: ALLRP-586022-23) awarded to DSM and a Principle Investigator Project Grant from the Austrian Science Fund (FWF, Grant Number PAT2648225) awarded to TH.

## AUTHOR CONTRIBUTIONS (https://credit.niso.org/)

CVS: Conceptualization, Formal Analysis, Methodology, Visualization, Writing – Original Draft, Writing – review & editing

CS: Conceptualization, Formal Analysis, Methodology, Writing – review & editing

DFC: Conceptualization, Methodology, Visualization, Writing – Original Draft, Writing – review & editing

DSM: Conceptualization, Formal Analysis, Methodology, Supervision, Writing – Original Draft, Writing – review & editing

TH: Conceptualization, Methodology, Project Administration, Resources, Supervision, Writing – Original Draft, Writing – review & editing

